# Relating the strength of density dependence and the spatial distribution of individuals

**DOI:** 10.1101/827238

**Authors:** Micah Brush, John Harte

## Abstract

Spatial patterns in ecology contain useful information about underlying mechanisms and processes. Although there are many summary statistics used to quantify these spatial patterns, there are far fewer models that directly link explicit ecological mechanisms to observed patterns easily derived from available data. We present a model of intraspecific spatial aggregation that quantitatively relates static spatial patterning to negative density dependence. Individuals are placed according to the colonization rule consistent with the Maximum Entropy Theory of Ecology (METE), and die with probability proportional to their abundance raised to a power *α*, a parameter indicating the degree of density dependence. Our model shows quantitatively and generally that increasing density dependence randomizes spatial patterning. *α* = 1 recovers the strongly aggregated METE distribution that is consistent with many ecosystems empirically, and as *α →* 2 our prediction approaches the binomial distribution consistent with random placement. In between our model predicts less aggregation than METE, but more than random placement. We additionally relate our mechanistic parameter *α* to the statistical aggregation parameter *k* in the negative binomial distribution, giving it an ecological interpretation in the context of density dependence. We use our model to analyze two contrasting datasets, a 50 ha tropical forest and a 64 m^2^ serpentine grassland plot. For each dataset, we infer *α* for individual species as well as a community *α* parameter. We find that *α* is generally larger in the tightly packed forest than the sparse grassland, and the degree of density dependence increases at smaller scales. These results are consistent with current understanding in both ecosystems, and we infer this underlying density dependence using only empirical spatial patterns. Our model can easily be applied to other datasets where spatially explicit data are available.

## INTRODUCTION

Quantitative understanding of spatial patterns in ecosystems can illuminate underlying processes (Levin, 1992; Rosenzweig, 1995; Brown et al., 2016) and allow us to better predict ecosystem response to natural and anthropogenic disturbances (Thomas et al., 2004; Newman et al., 2020). It is also critical in understanding the species-area relationship, a well studied (Arrhenius, 1921; Plotkin et al., 2000; Drakare et al., 2006; Harte and Kitzes, 2015) macroecological pattern, and has applications in reserve designs and conservation (Kitzes and Shirley, 2016).

Many different summary statistics have been used to quantify spatial patterning in ecology (Wiegand et al., 2013), and these summary statistics have been shown to be able to distinguish different ecological mechanisms (Brown et al., 2016). More broadly, there is evidence that emergent spatial patterns contain information about underlying ecological processes (Law et al., 2009; Brown et al., 2011; Münkemüller et al., 2020), meaning that spatial data can be used to infer the importance of various mechanisms. Rather than looking directly at summary statistics, we take a slightly different approach and directly model the impact of intraspecific negative density dependence on the spatially explicit species-level abundance distribution Π(*n|A, A*_0_*, n*_0_). This distribution is defined as the probability that if a species has *n*_0_ individuals in a plot of area *A*_0_, then it has *n* individuals in a randomly selected subplot of area *A*. We can then quantitatively examine how the degree of density dependence affects the shape of this distribution.

One prediction of this function comes from the Maximum Entropy Theory of Ecology (METE), which successfully and simultaneously predicts many macroecological patterns (Harte, 2011; Harte and Newman, 2014) across a wide range of spatial scales, taxa, and habitats (White et al., 2012; Xiao et al., 2015). For the Π(*n|A, A*_0_*, n*_0_) function, METE predicts very strong spatial aggregation consistent with many observed ecosystems. METE obtains the functional form of Π(*n*) by maximizing entropy while constraining the mean number of individuals in a subplot, however the same functional form can be obtained using a colonization rule.

Colonization rules assign spatial locations to new individuals based on the location of existing individuals. Harte (2011) shows that using the Laplace rule of succession as a colonization rule results in the same geometric distribution for Π(*n*) that METE predicts. Because METE agrees well with empirical data, we will use this colonization rule in our model. This is in contrast to a random colonization rule that would be consistent with the random placement model (Coleman, 1981). This model predicts no spatial aggregation and is therefore inconsistent with most observations, given that most species are aggregated (He and Gaston, 2000; Kitzes, 2019).

Empirically, we see that the degree of aggregation is sometimes less than the METE prediction (Conlisk et al., 2012). Conlisk et al. (2007) added an extra parameter to the relevant colonization rule that allows Π(*n*) to vary, but it has no mechanistic interpretation and is used only as a free fit parameter. Another approach is to fit the spatial data with a negative binomial distribution (Bliss and Fisher, 1953; He and Gaston, 2000, 2003; Green and Plotkin, 2007), but again this leaves us without any ecological intuition about the degree of aggregation to expect in different ecosystems as the additional parameter is a free fit parameter.

We derive a new model that uses the colonization rule consistent with METE and adds a density dependent death rule. Our model introduces a parameter *α* which quantifies the degree of intraspecific negative density dependence. Since this parameter has an ecological interpretation, we can use our model to infer the underlying density dependence from observed spatial patterning. This is in contrast to the models mentioned above, where the parameters have no mechanistic interpretation.

More generally, our model predicts a more random spatial arrangement with stronger negative density dependence and more spatial aggregation with weaker density dependence. While empirically there is an apparent qualitative relationship between species density and aggregation (Condit et al., 2000; Bagchi et al., 2011; Comita et al., 2014), our aim here is to establish a general quantitative statement relating density dependence and spatial aggregation.

First, we briefly review the METE and random placement predictions. In Methods, we derive our new probability distribution and explain how we will compare it to data, as well as introduce the datasets that we will be using for comparison. In Results, we look at the shape of this probability distribution and then present aggregate and species-level statistics for the data. Finally, in Discussion, we place our results for the analyzed datasets in a broader context, and describe possible applications.

## MATERIAL AND METHODS

### METE

In METE, the Π(*n*) function is given by (Harte et al., 2008)

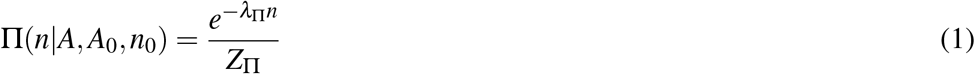

where *Z*_Π_ is a normalization factor, and *λ*_Π_ is the Lagrange multiplier determined by the constraint condition

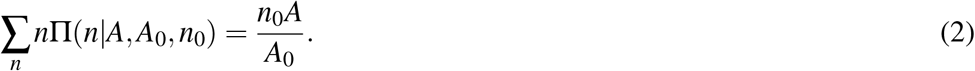

In the case of a bisection, *A* = *A*_0_*/*2 and the Π function simplifies to

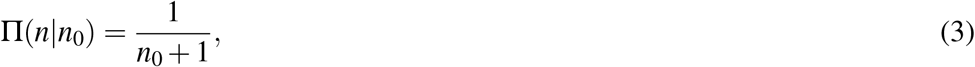

which is independent of *n*. This means that given *n*_0_ individuals, any arrangement of them on the two sides of a bisected plot or quadrat is just as likely as any other. Ecologically, this is equivalent to very strong spatial aggregation. This is in agreement with many datasets (Harte et al., 2008; Harte, 2011) but fails in others, where the theory over-predicts aggregation (Conlisk et al., 2007). This empirical agreement is why we choose the METE distribution as our starting point.

The prediction from METE is equivalent to the distribution obtained from using the Laplace rule of succession as a colonization rule (Harte, 2011). This rule states that in a colonization process, the probability of placing an individual on one side of the bisected area is roughly proportional to the fraction of individuals already there. This “rich get richer” effect results in strong spatial aggregation. More precisely, the probability for placing an individual on the left half of a bisected plot with *n*_*L*_ individuals on the left and *n*_*R*_ individuals on the right is

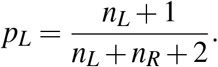

To make our notation consistent with that above, let the number on the left be *n* and the total number to be *n*_0_. The probabilities of a new individual arriving on the left or on the right are then:

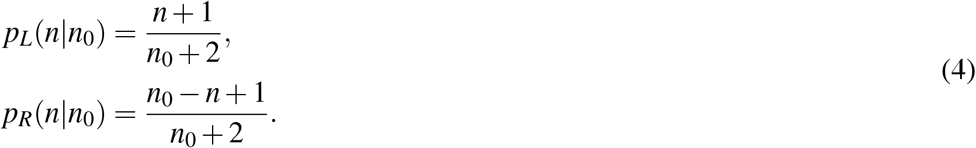

If we place *n*_0_ individuals using this rule, the resulting probability distribution is given by Eq. 3.

### Random placement

Another model for spatial ecology, perhaps the simplest, is the random placement model (Coleman, 1981). Instead of the placement rules in Eq. 4, each individual is placed randomly. In a bisected plot this means each individual has a 50 percent chance of being placed on either side, *p*_*L*_ = *p*_*R*_ = 0.5. Placing *n*_0_ individuals this way gives the binomial distribution

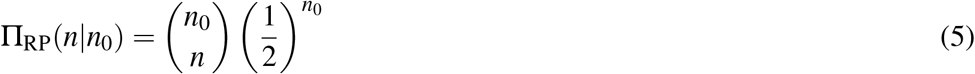

which, if *n*_0_ is large, means we are very likely to have roughly half the individuals on each side. This is equivalent to having no spatial aggregation; there is no preference for any new individual to stay close to any previous individual as each placement is a random coin flip.

### Introducing a density dependent death rule

We now introduce an intraspecific density dependent death rule in addition to the METE colonization rule in Eq. 4. To allow for general density dependence, we set the death rate proportional to *n*^*α*^. The parameter *α* determines the strength of the density dependence, and can be inferred from the data. Density dependence may result from resource limitation, or some other mechanism (e.g. the Janzen-Connell effect (Janzen, 1970; Connell, 1971)).

In the case of a bisected plot, each death must be on the left or right and the probability of a single death on the left, *p*_*D,L*_, or right, *p*_*D,R*_, is

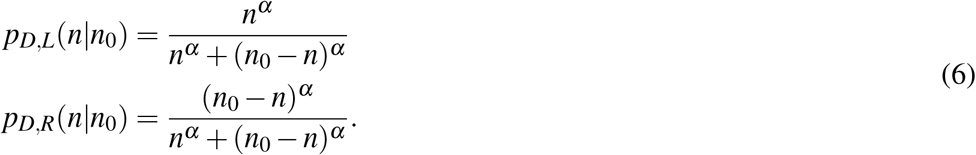

### Deriving the Π distribution

Now that we have the colonization and death rules (Eqs. 4 and 6), we can derive the general Π_*α*_(*n|n*_0_) for bisections. At each step in the model, we will have one death followed by the placement of one new individual. Each placement can be interpreted ecologically as a birth or as the immigration of an individual from the same species. This means the population size of the species is constant. Because we assume constant population size, we implicitly assume some degree of negative density dependence. However the strength of the density dependence is determined by the magnitude of *α*, where stronger density dependence corresponds to larger *α*. We can solve for the resulting steady state distribution where we reach an equilibrium in the spatial pattern.

There are several approaches for deriving the steady state solution for such a system. Here, we equate the rates leaving and entering any individual state Π_*α*_ (*n|n*_0_). We take the probability that we start with *n* individuals on the left, one on the right dies and then one is placed on the left resulting in *n* + 1 individuals on the left, and equate that to the probability that we have *n* + 1 individuals on the left, one on the left dies and then one is placed on the right resulting in *n* individuals on the left. We could have equivalently done the same thing with *n* and *n−* 1. Equating these rates using the probabilities in Eq. 4 and Eq. 6 leads to a recursion relation. Solving it gives a general stationary solution for Π_*α*_ with a given *n*_0_ and *α*:

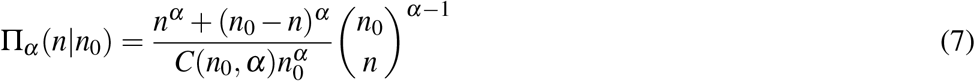

where *C*(*n*_0_*, α*) is the overall normalization that does not have a closed analytic form. In the case that *n*_0_(*α −* 1) is large, an approximate form for the normalization is

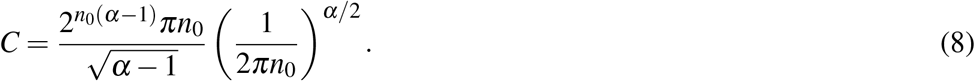

See Supporting information S1 for the details of this derivation.

If *α* = 2, we can solve for the normalization explicitly to get

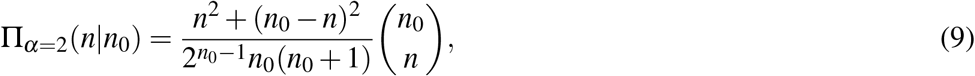

and if *α* = 1, we recover the METE prediction 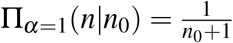.

### Beyond the first bisection

Bisecting the plot more than once (into quadrants, then into 8 cells, etc.) when comparing the model to data allows us to consider how aggregation changes with scale, as well as obtain multiple data points for individual species. We use the following method when we discuss bisecting a single plot more than once.

We begin by bisecting the plot in half in one direction, then bisecting each of the resulting plots in the opposite direction. We alternate this bisection pattern until we have 2^*b*^ cells, where *b* is the number of bisections. We can then compare adjacent cells as if they were their own bisection to obtain 2^*b−*1^ points.

When we go beyond the first bisection in the subsequent analysis, we only consider species that could have at least one individual per bisection (*n*_0_ *>* 2^*b−*1^). The smallest scale we consider in our datasets is *b* = 8, so when we bisect the plot more than once we will only consider species with more than 128 individuals. This restriction ensures that we do not have too many plots with only a few individuals present. If *n*_0_ is very small, Π_*α*_ (*n*) is not particularly sensitive to *α* and it becomes very difficult to reliably infer *α* from the data. For *n*_0_ *≤* 2, Π_*α*_ (*n*) does not depend on *α*.

### Inferring density dependence with α

Inferring the degree of intraspecific density dependence in empirical data requires obtaining a value of *α* consistent with the data. Bisection predictions can be compared to data by rank ordering the fraction of individuals present in one half of the plot for each species (e.g. Harte (2011)). This method, however, obscures trends in species abundance in the data and the predictions, and can lead to incorrect conclusions about which model is preferred (see Supporting information S2).

We instead find the maximum likelihood *α* given the data. Inferring *α* using this method gives us the values that are the most consistent with the data, even if they may not look like they agree with the rank ordered fractions (Supporting information Fig. S1 and Table S1).

For determining *α* for individual species, we will require multiple bisections and the sample size *p* will be roughly the number of cell pairings, *p ≈* 2^*b−*1^. There will be fewer data points in practice as some cell pairings will be empty.

We can also define a community *α*, assuming each species follows the same death rule with identical *α*. In this case, we will have a larger sample size. For a single bisection, we will have a sample size *p* equal to the number of species, *p* = *S*_0_. For multiple bisections where we consider the species on aggregate, the sample size will be roughly equal to the number of species multiplied by the number of cells, *p ≈ S*_0_2^*b−*1^. Again, the equality is not exact as not all cell pairings beyond the first bisection will have all of the species present at the single bisection level.

The statistical error in estimating *α* this way goes as 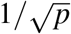 (see Supporting information S3). We can also get some idea of error from the maximum likelihood estimate itself by considering the width of the likelihood distribution, however for Fig. 3 and 4, we do not include these error bars as they are smaller than the data points.

### Data used

We will compare our results to two contrasting datasets. First, we will use data from a sparse Californian serpentine grassland site (Green et al., 2003) at the McLaughlin University of California Natural Reserve censused in 1998. This is a 64 m^2^ plot divided into 256 cells with 24 species and 37 182 individuals. There are 10 species with abundance greater than 128 individuals that constitute 36 783 individuals. The serpentine data are available from the Dryad Digital Repository at https://doi.org/10.6078/D1MQ2V Green et al. (2019).

Second, we will use data from Barro Colorado Island (BCI) in Panama (Condit, 1998; Hubbell et al., 1999, 2005), a 50 ha plot in a moist tropical forest. We will work with the 2005 census and consider plants with a diameter at breast height (dbh) greater than 10 cm. This dataset has 229 species and 20 852 individuals, and 40 species with abundance greater than 128 individuals that constitude 15 960 individuals. The BCI data can be found at https://forestgeo.si.edu/explore-data and are available from the Dryad Digital Repository at https://doi.org/10.15146/5xcp-0d46 Condit et al. (2019).

## RESULTS

### Comparison to METE and random placement

Figure 1 compares the bisection predictions for Π(*n*) from METE, random placement, and our density dependent model for various *α*, at *n*_0_ = 10 and 50. In general, our model predicts that increasing negative density dependence (larger *α*) leads to more random spatial patterning, and less density dependence (smaller *α*) leads to stronger aggregation.

**Figure 1.**
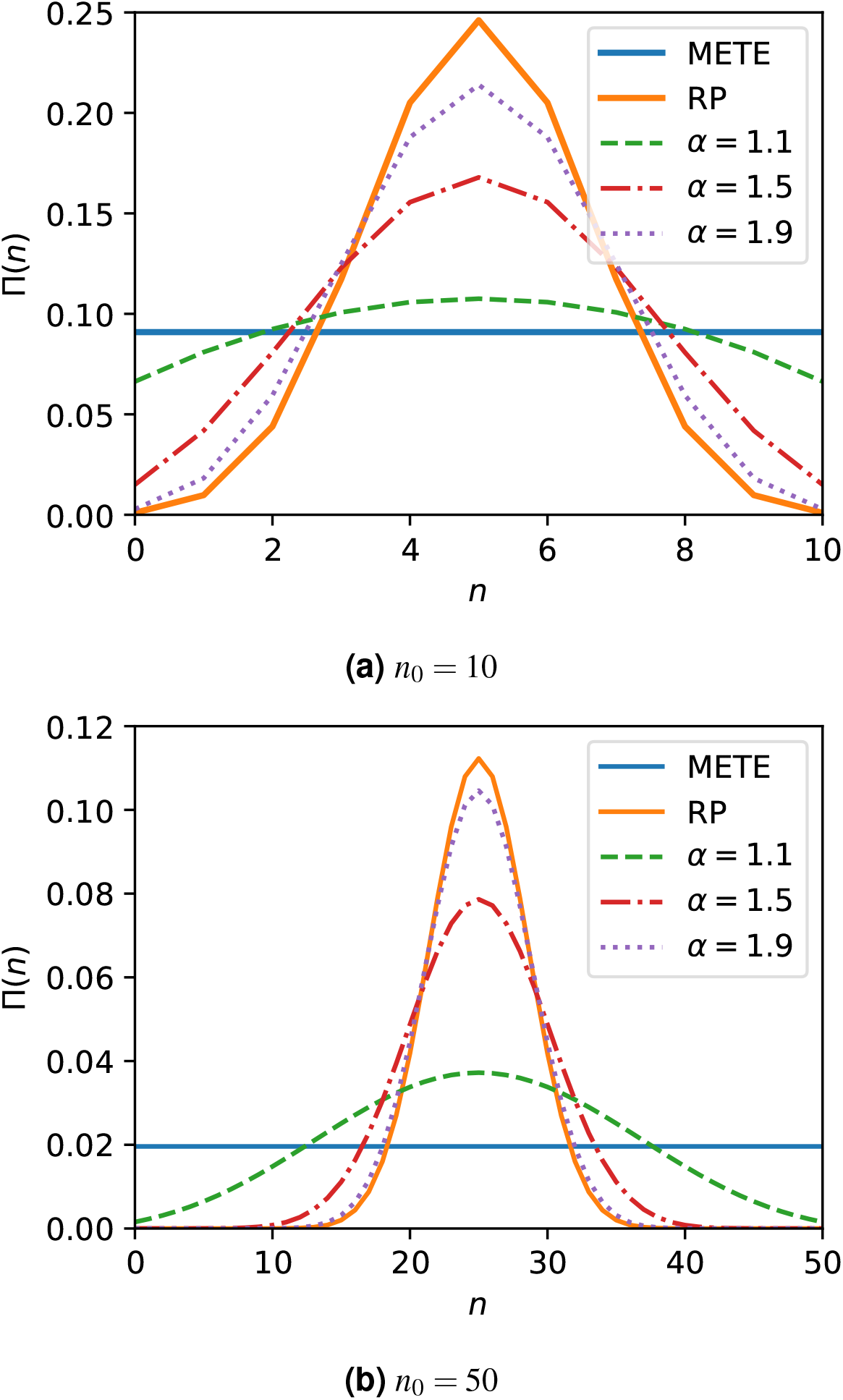
Comparison of the bisection probability distributions Π(*n*) from METE, random placement (RP), and our density dependent model with varying *α*. At *α* = 1, our model corresponds exactly to METE. At larger *n*_0_, *α* → 2 approaches the random placement distribution. Our model varies continuously between METE and random placement for 1 *< α <* 2.

We can relate our distribution directly to both the METE and random placement distributions for different values of *α*. *α* = 1 corresponds exactly to the METE solution, which makes sense given that the placement and death rules are both linear in *n*. As *α →* 2, our distribution approaches the random placement prediction if *n*_0_ is large enough (Supporting information S4 shows this result analytically). For 1 *< α <* 2, we vary continuously between METE and random placement. We can make the distribution even more spatially aggregated than METE with *α <* 1 and even less than random placement (overdispersed) with *α >* 2.

We can also relate this distribution to the commonly used conditional negative binomial distribution (Bliss and Fisher, 1953; He and Gaston, 2000, 2003; Green and Plotkin, 2007) in the limit of large *n*_0_, assuming that matching the peak of the distributions is a good approximation for the entire distribution. In that limit, the aggregation parameter *k* is approximately related to the density dependent parameter *α* by

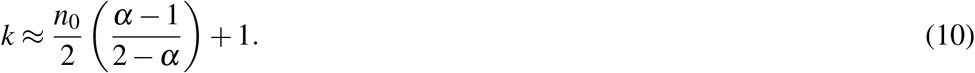

Note that this approximation holds for 1 *≤ α ≤* 2, which is the ecologically relevant range for most species. See Supporting information S5 for the derivation.

### Individual species

Since the Π function is defined on the species level, we can consider each species separately and find the maximum likelihood *α* for each. To do this we have to go beyond the first bisection to get multiple data points for the same species at smaller scales.

For the serpentine data, we exclude *Eriogonum nudum* from the following figures as an outlier (see Discussion). This leaves 9 species with abundance greater than 128 individuals.

Figure 2 shows the distribution of *α* values among the species at each scale, for both the serpentine and BCI data. The median *α* increases at smaller scales for both datasets, and is higher overall at the BCI dataset, even though the absolute scale is much larger. The spread in *α* is quite large, but this variation is expected considering the small number of data points, especially for rarer species. Most species have an *α* between 1 and 2, which is somewhere between the aggregation predicted by METE and random placement.

**Figure 2.**
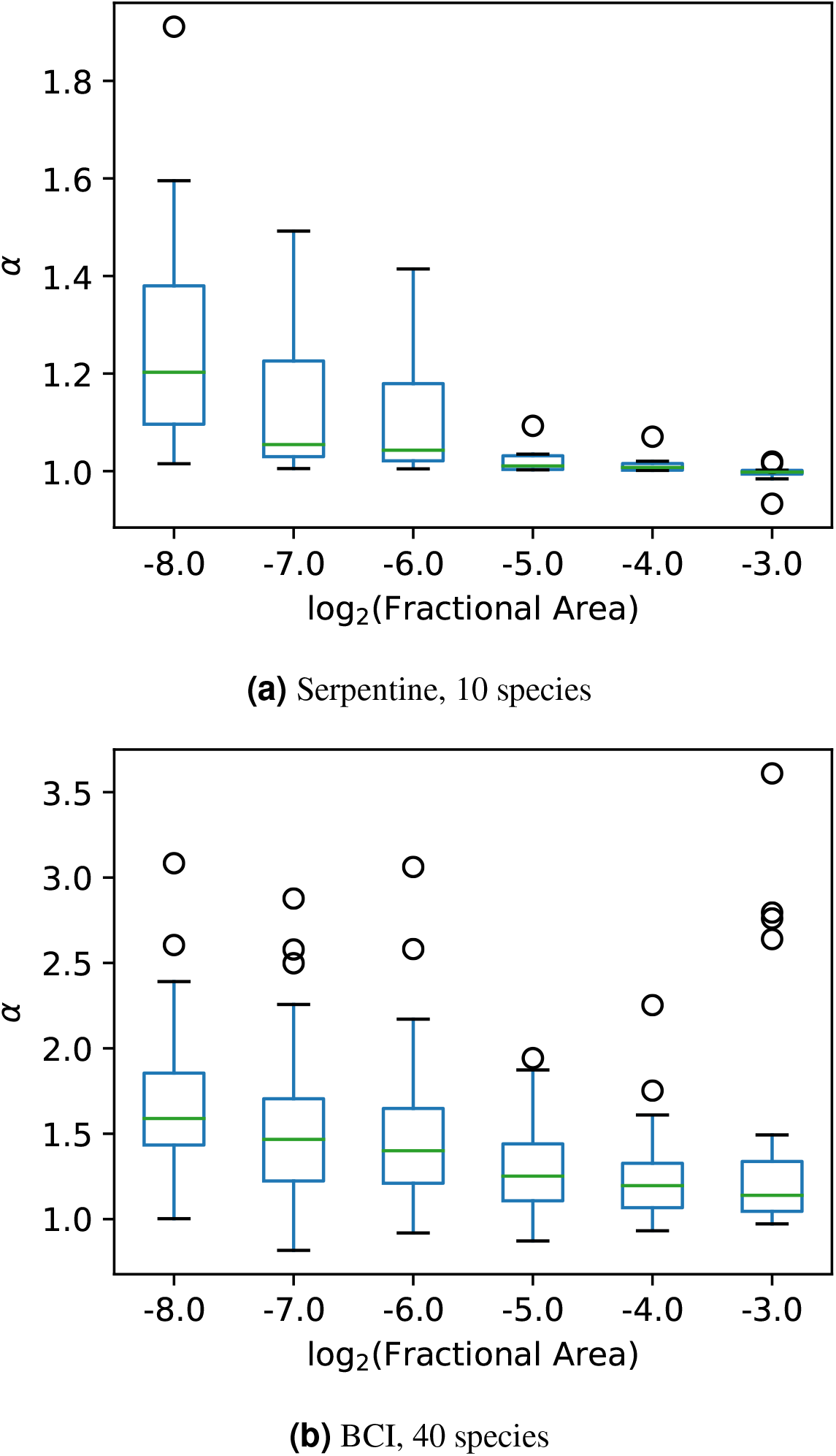
Boxplots for *α* among the species at different scales. At smaller scales *α* is larger, and we see a relatively large spread in *α* across species at the same scale. The boxplots show boxes from quartile 1 (Q1) to quartile 3 (Q3) with a line at the median. The whiskers extend to 1.5×(Q3-Q1). The remaining points are plotted as individual circles.

### Community α

We can instead treat *α* as a community parameter, using each species as a single data point to recover a community *α*. Figure 3 shows the direct comparison between our model and the serpentine and BCI datasets at the single bisection level. Each data point is the observed fraction of individuals in one half of the plot versus the species abundance. The curves in this figure show the 95% contour intervals for Π(*n*) distributions predicted by METE, random placement, and our density dependent model with the maximum likelihood *α* value. The contours narrow at larger *n*_0_ as the distributions narrow.

**Figure 3.**
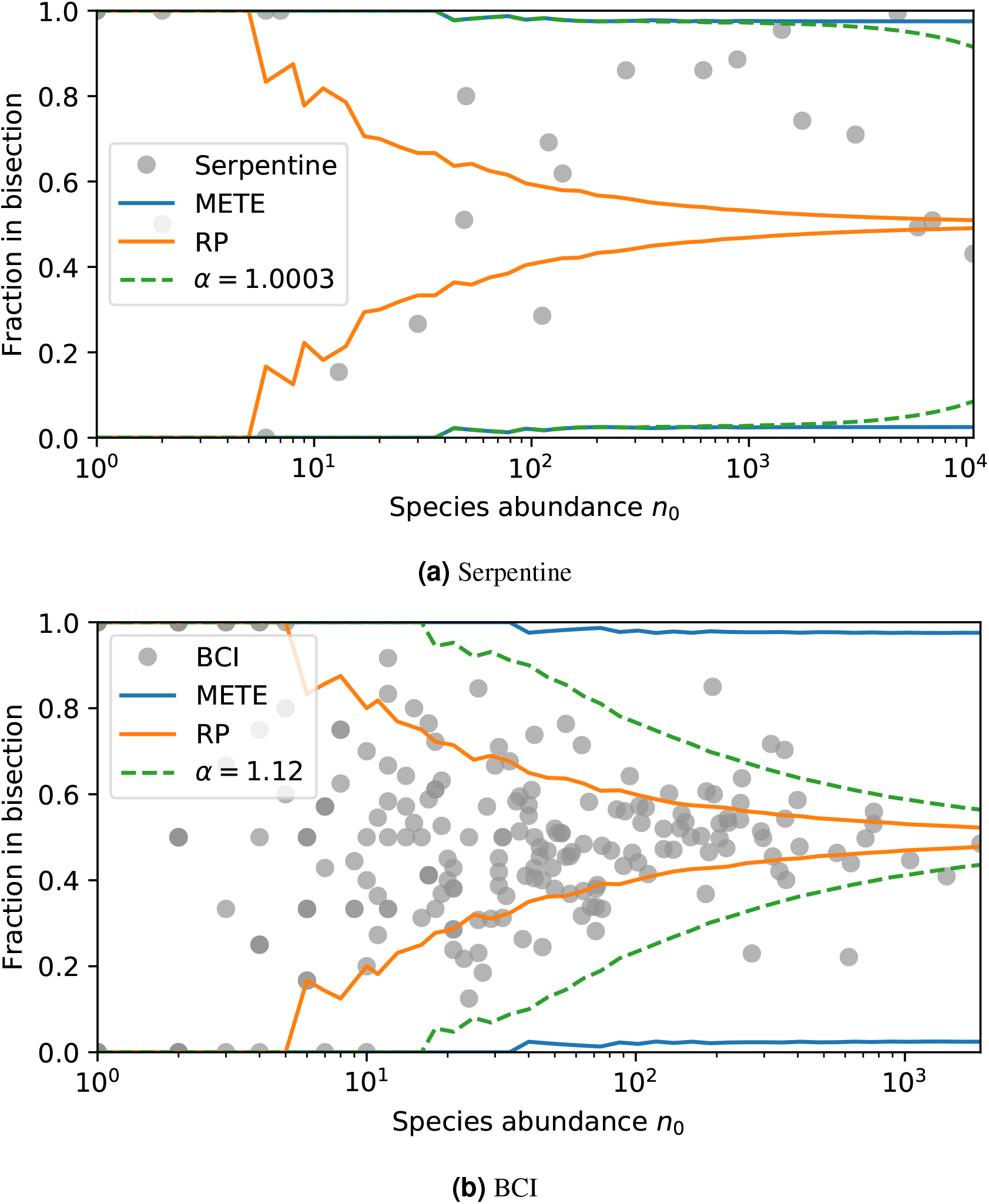
95% contour intervals for the different predicted probability distributions overlayed with (3a) the serpentine dataset, and (3b) the BCI dataset. For our density dependent model with a community *α*, *α* = 1.0003 maximizes the log-likelihood for the serpentine dataset, and *α* = 1.12 maximizes log-likelihood for BCI dataset.

At the single bisection level, the maximum likelihood result for the serpentine dataset is nearly indistinguishable from *α* = 1, so the confidence interval curves on the plot for METE and the density dependent model overlap for most *n*_0_. For the BCI data, the maximum likelihood value is *α* = 1.12, slightly larger than 1. We can see the contours in this case are clearly in between those for METE and random placement. The likelihoods for each of the models are shown in Table 1.

**Table 1.** Log-likelihood values for the three different models, with *α* as a community parameter. We can compare our model to METE using the deviance in log-likelihood and obtain a *p*-value. The deviance is defined as twice the difference in log-likelihood. For the serpentine dataset, the deviance is 0.6 which corresponds to a *p*-value of 0.45. For the BCI dataset, the deviance is 138 which corresponds to a *p*-value of *<* 10^*−*30^.

| Serpentine |  | BCI |  |
| --- | --- | --- | --- |
| Model | Log-likelihood | Model | Log-likelihood |
| METE | -114.8 | METE | -729 |
| RP | -5188.6 | RP | -963 |
| $\alpha = 1.0003$ | -114.6 | $\alpha = 1.12$ | -660 |

### Scale dependence in community α

Going beyond the first bisection allows us to see how *α* varies depending on the scale of our plot. Figure 4 shows how *α* scales with fractional area for both the serpentine and BCI plots. Density dependence increases at smaller scales in both datasets. The trend in community *α* across scales is similar to the median *α* in the single species analysis, though the median *α* is in general slightly larger than the community *α*. Note that here we restrict our analysis to species with *n*_0_ *>* 128 for all scales so that we are including the same species across scales.

**Figure 4.**
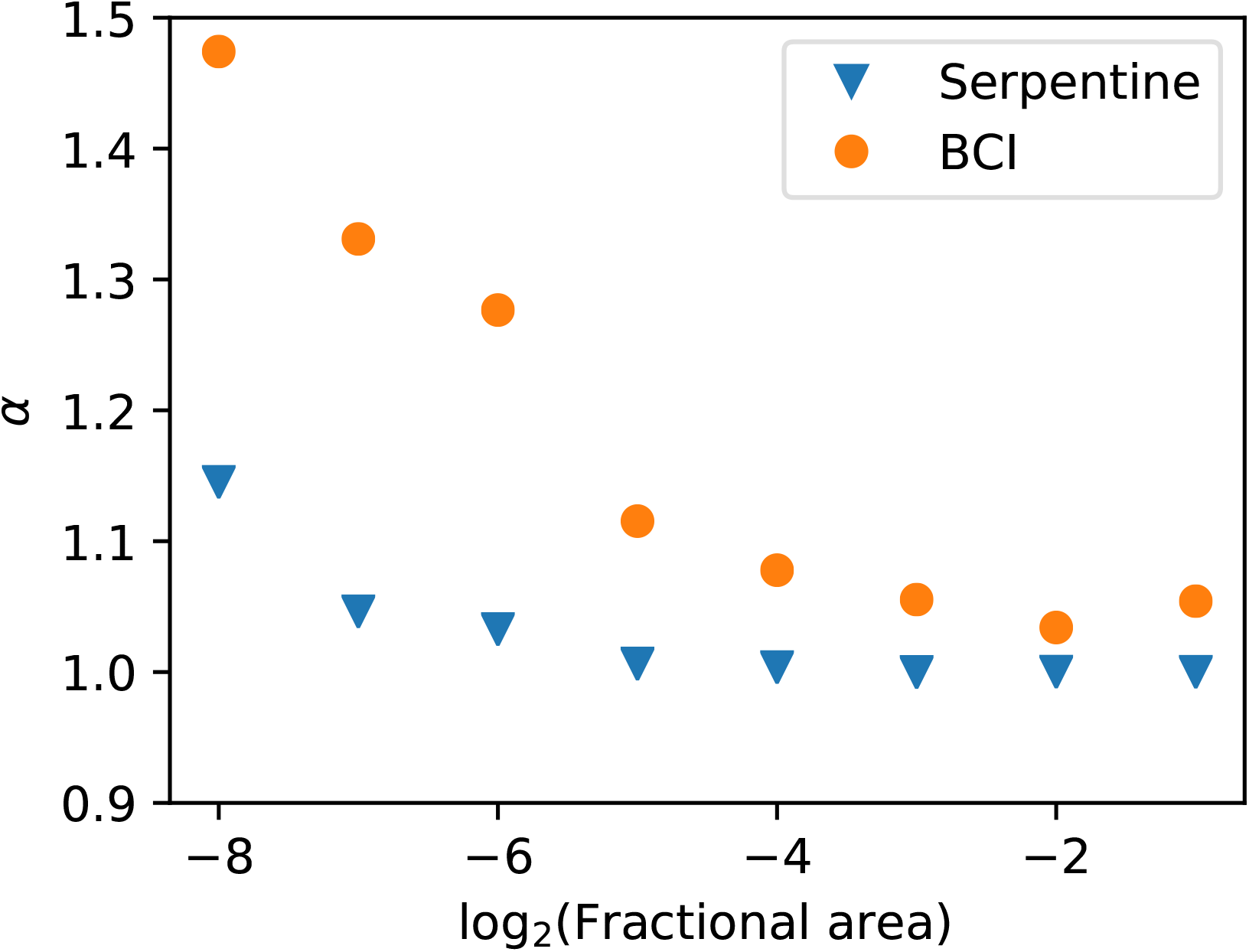
Community *α* scaling with area for species with abundance *n*_0_ *>* 128. The density dependence again increases at smaller scales and the trend is similar to the single species analysis. The serpentine dataset has 36 783 individuals and the BCI dataset has 15 960 individuals.

## DISCUSSION

Our model establishes a quantitative relationship between the spatially explicit distribution Π(*n*) and the parameter *α*, which measures the strength of negative density dependence. This can be seen in Fig. 1, where the Π_*α*_ (*n*) distribution flattens with smaller *α*, indicating greater aggregation, and broadens as *α* increases. Importantly, the parameter *α* has a direct interpretation. Unlike the models outlined in the introduction (Bliss and Fisher, 1953; He and Gaston, 2000, 2003; Green and Plotkin, 2007; Conlisk et al., 2007) where an ad hoc parameter is tuned to fit spatial patterns, our model introduces a parameter that models the strength of negative density dependence. Further, our relationship in Eq. 10 allows us to interpret the parameter *k* in the negative binomial distribution in the same intuitive way.

### Comparing species and community α

In our analysis, we consider *α* both as a separate parameter for each species (as in Fig. 2, and as a community parameter where each species has the same *α* (as in Fig. 3 and Fig. 4). The community *α* is harder to interpret ecologically, but we include it to allow for comparisons with models with community level aggregation parameters (e.g. Conlisk et al. (2007) and Volkov et al. (2005)). To analyze and compare the accuracy of the species-level *α* and the community *α*, we considered the Akaike Information Criterion (AIC) in both cases across scales (Table 2). For both serpentine and BCI at all scales considered, we find that the AIC is lower with species-level *α* compared to a single community *α*, despite the inclusion of 9 more parameters in the case of the serpentine data and 228 more parameters in the case of the BCI data. This leads us to believe that a separate *α* for each species describes the data better than a single community *α*.

**Table 2.** Comparison of the Akaike Information Criterion (AIC) for *α* defined at the individual species level and at the community level in both the serpentine and BCI data and across scales. The AIC is lower at the species level in all cases.

| **Scale ($A/A_0$)** | | $2^{-8}$ | $2^{-7}$ | $2^{-6}$ | $2^{-5}$ | $2^{-4}$ | $2^{-3}$ |
| --- | --- | --- | --- | --- | --- | --- | --- |
| **Serpentine** | Species $\alpha$, AIC | 474 | 769 | 1294 | 1931 | 3208 | 4881 |
| | Community $\alpha$, AIC | 485 | 777 | 1321 | 2079 | 3420 | 5182 |
| **BCI** | Species $\alpha$, AIC | 10133 | 7541 | 5079 | 3409 | 2109 | 1271 |
| | Community $\alpha$, AIC | 10207 | 7621 | 5148 | 3522 | 2171 | 1307 |

### Comparing serpentine and BCI

We use our model to directly compare our results between our two contrasting datasets, serpentine and BCI. Because the serpentine site was very sparse, whereas the BCI forest is tightly packed, we expect higher *α* values and greater density dependence at BCI than at the serpentine site. This is consistent with our inferred values of *α* at both the individual species level and at the community level.

Another difference between the serpentine and BCI sites is how well other macroecological distributions agree with METE. METE well describes other patterns at the serpentine site, and does less well at explaining the BCI data. Given that *α* = 1 corresponds to the METE prediction for Π(*n*), we might expect that ecosystems well described by other METE predictions will have *α ≈* 1, as these systems will generally be consistent with METE. This could be the case as METE predictions seem to hold for relatively static and undisturbed ecosystems (Newman et al., 2020). This interpretation is consistent with our analysis here as the median and community *α*s for the serpentine data are approximately 1 at the largest scale, whereas at BCI the median and community *α*s are larger than 1.

### Scaling

Our scaling results in both Figs. 2 and. 4 make ecological sense. We expect that at smaller scales, the density dependence would be larger as individuals compete more for resources at that scale. At large scales, we expect *α* to be close to 1 as the individuals do not compete over large distances. This means that the spatial distributions look more aggregated on large scales than on small scales as the individuals within species broadly group together, but repel each other at small scales. We see this trend at the individual level in Fig. 2 as the medians increase at smaller scales, and for the community *α* in Fig. 4.

Our repeated bisection analysis also indicates at which scale density dependence becomes important. This will appear as a shoulder in the data where *α* moves away from *≈* 1. We could do this for individual species by tracing *α* and looking for a shoulder in Fig. 2, but here we will look at the community results in order to compare to Conlisk et al. (2007) and Volkov et al. (2005). Looking at Fig. 4, we find that the shoulder in absolute scale corresponds to *<* 0.5m^2^ for the serpentine plot and *<* 1.6 ha for the BCI dataset. This again makes sense given that the serpentine grassland is much more sparse than the BCI forest.

We first compare to Conlisk et al. (2007), who introduce a fit parameter *ϕ* that modifies the colonization rule Eq. 4 and allows the Π distribution to vary continuously between random placement and METE. For the serpentine data, they find that at scales larger than around 0.5 m^2^ (the 8th bisection), *ϕ* approaches 0.5, which corresponds to the METE prediction. At scales around 0.5 m^2^ or smaller, *ϕ ≈* 0.2, where *ϕ* = 0 corresponds to random placement. This is consistent with our scaling results in Fig. 4.

Volkov et al. (2005) showed that intraspecfic and symmetric density dependence can explain different shapes for the species-abundance distribution. Their added parameter *c* is interpreted as a measure of the strength of symmetric density dependence, where *c* = 0 corresponds to no density dependence. This parameter is therefore similar to our community *α* in that all species have the same degree of negative density dependence. They then show at what density these effects become important in their Fig. 3. For BCI, they find *c* = 1.80, and the density dependent effects are visible for species with *n >* 27. This corresponds to density dependence entering at scales smaller than a fractional area of 1*/*27 = 1*/*2^4.75^, which is close to the same scale where we see *α* increase away from 1 in Fig. 4.

At the individual species level at BCI, Condit et al. (2000) find that individuals are clustered at small scales and distributed randomly at large scales. Though initially this result seems to be in conflict with our results, it is explained by a different measure of aggregation. They use Ω_*x*_, which is the total number individuals found within a radius *x* of a single individual, normalized by the average density of individuals. Ω_*x*_ = 1 corresponds to random placement, and Ω_*x*_ > 1 indicates aggregation. At small scales at BCI, they find Ω_0*−*10m_ *≫* 1, and at large scales Ω_*>*100m_ *≈* 1. We find that for most species, *α >* 1 at small scales and *α ≈* 1 at large scales (Fig. 2). To see that these results are consistent, consider a plot with a cluster of individuals on one side where the spatial structure within the cluster is random. At the single bisection level this would be consistent with *α* = 1, but within the cluster *α >* 1. That same arrangement of individuals would give an Ω_*x*_ that shows strong aggregation at small scales and random placement at large scales. This is what we find empirically in the BCI data. This spatial pattern is also consistent with the Poisson cluster model outlined in Plotkin et al. (2000), where clusters of individuals are placed uniformly with random internal spatial structure.

### Trends for individual species at BCI

At the individual species level more broadly, we might expect to find trends in inferred density dependence with species abundance or size. More abundant species may be competing more for the same resources, or larger species may compete over larger distances. For example, Condit et al. (2000) find that both rarer species and smaller individuals tend to be more aggregated, however at a much smaller scale (within a 10 m radius) and with a different metric of aggregation.

In the BCI data, we do not find any species with high *α* and high abundance (Supporting information Fig. S4), but otherwise we do not see a trend in abundance. We also find no evidence of a trend with species’ mean dbh (Supporting information Fig. S5), though it is possible this trend is obscured by variance in individual size within a species, or that the range of mean dbh we considered (about 100 *−* 500 mm) is too small to see its effect.

Finally, we looked for an overall energy effect. Considering that the most abundant species tend to be smaller, it may be that density dependence depends on the total metabolic rate of a species. Plotting this relationship (Supporting information Fig. S6) again does not reveal a scaling relationship.

That none of these trends are found in the data might indicate that the operating density dependent mechanism is not resource limitation, in that it doesn’t depend on the abundance, size, or metabolic rate of the species. A plausible mechanism for the observed density dependence at BCI is the Janzen-Connell effect (Janzen, 1970; Connell, 1971), whereby areas near parent trees are inhospitable for offspring, resulting in density dependence. Various studies (Harms et al., 2000; Carson et al., 2008; Comita et al., 2014) have observed this effect at BCI, which is consistent with our result that *α >* 1 for most species at smaller scales there. See Supporting information S6 for more information on these trends.

### Notable species

For individual species at the BCI dataset, *Gustavia superba* stood out with an average *α* of 1.001 across scales. This species is largely limited to 2 hectares of young secondary forest along the edge of the plot, (J. Wright, personal communication, 2019) making it look especially aggregated and resulting in a maximum likelihood *α* close to 1.

In the serpentine dataset, we excluded *Eriogonum nudum* as an outlier for part of our analysis. The maximum likelihood *α* was *>* 6 at the smallest scale and the maximum itself was very shallow. This species has a large canopy compared to the other grassland plants, and tends to be found far from other individuals. It makes sense that it would be overdispersed with *α >* 2.

### Implications for the species-area relationship

In METE, the spatial distribution is used together with the species-abundance distribution to predict the species-area relationship (Harte, 2011), and to upscale predictions of biodiversity (Harte and Kitzes, 2015). These predictions should hold in ecosystems like the serpentine grassland analyzed here, as the observed species aggregation agrees with the METE prediction. However, different levels of aggregation will impact the species-area relationship. The impact of aggregation is discussed in Wilber et al. (2015). They find that increasing randomization decreases the predicted slope of the species-area relationship at the same scale, and therefore upscaling METE will overpredict species richness. In addition, they analyze the effect of variation in aggregation among species, which slightly decreases the slope at small scales and increases the slope at larger scales. This results in a species-area relationship that more closely resembles a power law. They also consider the effect of decreasing aggregation across scale, which results in a species-area relationship that no longer displays scale collapse. We observe both of these effects here.

### Limitations and assumptions

Our model considers only bisections, and it would be useful to extend it to be more general. There are many spatial arrangements that can not be accurately captured by dividing plots into bisection, and in general a single functional summary statistic does not completely describe the observed spatial pattern (Wiegand et al., 2013). For example, if we divide our plot into an *m* by *m* grid, and have one individual per cell, we would see exactly 0.5 as the fraction for each bisection. This result would be consistent with random placement with a large number of individuals, which does not well describe this exceptionally uniform arrangement. There could also be different degrees of spatial aggregation within a cell that we will not accurately capture with a bisection. Despite these limitations, bisections are useful for understanding commonly observed macroscopic spatial patterns.

A conceptually simple extension to our model is to divide plots into quadrisections rather than bisections. The colonization and death rules then have three unknowns rather than one (the number of individuals in each quadrant, where the fourth is determined by constraining the sum to be *n*_0_). This makes it hard to solve analytically, however we can simulate the birth-death process until it reaches steady state. We find no significant difference in our simulation compared to our prediction from two bisections, and find that a community *α* = 1.12 is still consistent with the BCI data.

Because we consider the steady state solution in our model, we are assuming that the density dependence time scale is longer than the time scale of individual births or deaths. That is, *α* must not change too rapidly in time. This assumption is justified for many systems roughly in steady state with their environment and not undergoing rapid change (Newman et al., 2020).

Solving for the steady state solution also assumes that births and deaths are in balance. We assume here that there is a single death followed by a placement, however simulating two deaths followed by two placements gives a probability distribution consistent with our analytic prediction. We expect our result to hold with other numbers of deaths and placements. Assuming that births and deaths are in balance also implicitly assumes some amount of negative density dependence, and here *α* provides a quantitative measure of the degree of density dependence.

Another assumption in our model is the choice of colonization rule itself, though if we had chosen a different colonization rule many of our conclusions would remain the same. We use the colonization rule consistent with METE because of its good empirical agreement (Harte, 2011). This allows us to interpret the *α* = 1 case as consistent with METE. This is useful as METE can be thought of as a null theory that holds in ecosystems that are undisturbed and relatively static (White et al., 2012; Xiao et al., 2015; Harte et al., 2017; Newman et al., 2020), and *α* ≠ 1 can be thought of as a density dependent correction.

As an example, we could have chosen the colonization rule resulting in the random placement distribution, which for a bisection is just *p*_*L*_ = *p*_*R*_ = 0.5. In this case, *α* = 1 would recover the binomial distribution which we know does not well describe most spatial data (He and Gaston, 2000), and so we cannot interpret *α* ≠ 1 as a density dependent correction. As another example, if we had chosen the more general colonization rule in (Conlisk et al., 2007) we would have two parameters to tune, making it difficulty to differentiate between colonization and death. In ecosystems where we suspect a different colonization rule may be in play, we could modify our theory appropriately. In any of these cases, our general results would remain largely unchanged.

## Conclusion

Our model robustly predicts that increased intraspecific negative density dependence leads to more random spatial patterning, and establishes a quantitative relationship between the degree of density dependence described by the parameter *α* and spatial patterning described by the metric Π(*n*). We predict that this result is general across ecosystems and taxonomic groups. We find that at all but the smallest scales, the serpentine grassland site is consistent with the absence of a density dependent correction and has the strong spatial aggregation predicted by METE. This is true for both the median individual species and at the community level. At the tropical forest site, negative density dependence is important: the median species *α* and the community *α* are both greater than 1 at even the largest scales. Both ecosystems show scaling of negative density dependence as we expect with median species *α* and community *α* larger at smaller scales, and increasing away from 1 at scales consistent with other analyses. Overall, our results are consistent with current understanding at both sites. Our model allows us to infer this underlying density dependence using only static spatial patterning, and can be applied in any ecosystem with spatially explicit data.

## Data availability statement

The serpentine data are available from the Dryad Digital Repository at https://doi.org/10.6078/D1MQ2V. The BCI data can be found at https://forestgeo.si.edu/explore-data and are available from the Dryad Digital Repository at https://doi.org/10.15146/5xcp-0d46. Relevant code is available at https://github.com/micbru/density_dependence_public.

## Declarations

## Acknowledgements

JH thanks the Santa Fe Institute for their hospitality. We thank Kaito Umemura for valuable discussion and feedback, and Egbert Leigh and Joseph Wright for their help with the BCI dataset. We thank Jessica Green for the serpentine data. The BCI forest dynamics research project was founded by S.P. Hubbell and R.B. Foster and is now managed by R. Condit, S. Lao, and R. Perez under the Center for Tropical Forest Science and the Smithsonian Tropical Research in Panama. Numerous organizations have provided funding, principally the U.S. National Science Foundation, and hundreds of field workers have contributed.

## Author contributions

Both authors were involved in conceptualization. MB conducted the formal analysis and led the writing of the manuscript.

