## Supporting Information for "Relating the strength of density dependence and the spatial distribution of individuals"

#### CONTENTS

**S1 Derivation of the probability distribution** **2**

**S2 The problem with rank ordered fractions** **4**

**S3 Error in recovering $\alpha$** **8**

**S4 Relating to the binomial distribution** **10**

**S5 Relating to the negative binomial distribution** **11**

**S6 Trends in diameter and abundance for BCI data** **14**

**References** **17**

#### S1 DERIVATION OF THE PROBABILITY DISTRIBUTION

Following the methods in the main text and equating the rates entering and exiting $\Pi(n)$ gives

$$p_R(n|n_0-1)p_{D,L}(n+1|n_0)\Pi(n+1|n_0) = p_L(n|n_0-1)p_{D,R}(n|n_0)\Pi(n|n_0).$$

Plugging in with Eq. 4 and 6 gives

$$\frac{(n+1)^{\alpha-1}}{(n_0-n-1)^{\alpha}+(n+1)^{\alpha}}\Pi_\alpha(n+1) = \frac{(n_0-n)^{\alpha-1}}{(n_0-n)^{\alpha}+n^{\alpha}}\Pi_\alpha(n). \tag{S1}$$

We can solve this recursion relation to obtain a general stationary solution. To do so, we define

$$P(n) = \frac{\Pi(n)}{(n_0-n)^{\alpha}+n^{\alpha}}$$

to get

$$(n+1)^{\alpha-1}P(n+1) = (n_0-n)^{\alpha-1}P(n).$$

We can fix $\Pi(0)$, since we can choose it freely before normalizing the distribution. For simplicity, choose $\Pi(0) = 1$. Then we can write the first few $P$ terms and generalize to $P(n)$:

$$\begin{aligned}
P(0) &= \frac{1}{n_0^{\alpha}}\\
P(1) &= \frac{1}{n_0}\\
P(2) &= \left(\frac{n_0-1}{2}\right)^{\alpha-1}\frac{1}{n_0}\\
P(3) &= \left(\frac{(n_0-1)(n_0-2)}{6}\right)^{\alpha-1}\frac{1}{n_0}\\
&\vdots\\
P(n) &= \left(\frac{(n_0-1)(n_0-2)\dots(n_0-(n-1))}{n!}\right)^{\alpha-1}\frac{1}{n_0}\\
&= \left(\frac{n_0!}{n!(n_0-n)!}\right)^{\alpha-1}\frac{1}{n_0^{\alpha}}.
\end{aligned}$$

Finally, we plug this back into our definition of $\Pi(n)$ to obtain the general stationary $\Pi(n)$ for a given $n_0$ and $\alpha$

$$\Pi_\alpha(n|n_0) = \frac{n^{\alpha}+(n_0-n)^{\alpha}}{C(n_0,\alpha)n_0^{\alpha}}\binom{n_0}{n}^{\alpha-1} \tag{S2}$$

which is Eq. 7 in the main text.

We now want an approximate analytic form for the normalization $C(n_0, \alpha)$,

$$C(n_0,\alpha) = \sum \frac{(n^\alpha + (n_0-n)^\alpha)}{n_0^\alpha} \binom{n_0}{n}^{\alpha-1}. \tag{S3}$$

We begin by approximating the binomial term with a normal distribution, which is good for large $n_0$. This gives

$$\binom{n_0}{n}^{\alpha-1} \approx \exp\left(-\frac{2}{n_0}(\alpha-1)(n-n_0/2)^2\right)\left(2^{-n_0}\sqrt{\frac{\pi n_0}{2}}\right)^{-(\alpha-1)}.$$

We then use the symmetry of the sum over $n$ to write $n^\alpha + (n_0-n)^\alpha = 2n^\alpha$. Finally, we replace the sum with an integral over $n$ from 0 to infinity, again assuming $n_0$ is very large. This integral gives

$$\int_0^\infty dn 2n^a \exp\left(-\frac{2}{n_0}(\alpha-1)(n-n_0/2)^2\right) =$$
$$\left(\frac{n_0}{2(\alpha-1)}\right)^{\alpha/2}\left(\frac{\alpha n_0}{2}\Gamma\left(\frac{\alpha}{2}\right){}_1F_1\left(\frac{1-\alpha}{2},\frac{3}{2},-\frac{n_0}{2}(\alpha-1)\right)+\right.$$
$$\left.\sqrt{\frac{n_0}{2(\alpha-1)}}\Gamma\left(\frac{\alpha+1}{2}\right){}_1F_1\left(-\frac{\alpha}{2},\frac{1}{2},-\frac{n_0}{2}(\alpha-1)\right)\right). \tag{S4}$$

Now if we assume $n_0(\alpha-1)$ is large, we can approximate

$${}_1F_1(a,b,z) \approx \frac{\Gamma(b)}{\Gamma(b-a)}(-z)^{-a}$$

and Eq. S4 becomes

$$2\sqrt{\frac{\pi}{\alpha-1}}\left(\frac{n_0}{2}\right)^{\alpha+1/2}.$$

Putting the other contributions to the normalization back in, we get

$$C(n_0,\alpha) = \frac{2^{n_0(\alpha-1)}\pi n_0}{\sqrt{\alpha-1}}\left(\frac{1}{2\pi n_0}\right)^{\alpha/2}, \tag{S5}$$

as Eq. 8 in the main text.

#### S2 THE PROBLEM WITH RANK ORDERED FRACTIONS

As mentioned in the main text, Harte (2011) and others have compared bisection predictions to data by rank ordering the fraction of individuals present in half of the plot for each species. This gives a number of points equal to the number of species in the plot. Rank ordering in this way with the BCI data gives us 229 data points, one for each species.

Since our models only predicts the bisection curve for a single $n_0$, but our data incorporates species with different $n_0$, we still have to do a bit of work to compare our theory to the rank ordered data. For each species, we generate one value of $n$ randomly from the desired theoretical distribution with abundance $n_0$. We then rank order these predicted fractions and compare to the rank ordered data. Figure S1a shows the results using the BCI data.

It looks like $\alpha = 1.4$ is a very good fit to the data, which we obtained by eye to the first decimal place. With this method, the BCI data deviated the most from the METE prediction for the datasets considered in Harte (2011).

To fit the free parameter $\alpha$ more rigorously, we maximize the log-likelihood of the density dependent distribution given $n$ and $n_0$ from the data. We obtain $\alpha = 1.12$, which would not appear to be a good fit to the data if we had rank ordered.

The log-likelihood values from the rank-ordered best fit $\alpha$, the maximum log-likelihood $\alpha$, as well as random placement, and METE are given in Table S1. Note that even though our model with $\alpha = 1.4$ looks like it has a much better fit in Fig. S1a, the log likelihood is very close to that of METE, whereas $\alpha = 1.12$ provides a much better fit to the data.

Unfortunately, rank ordering the results from the distribution draws in Fig. S1a hides how likely specific points are. There are some points in Fig. S1a that are actually very unlikely, but we don't see them in the rank ordered plot. This is because the probability distribution changes with different $n_0$, but Fig. S1a puts all the points together in one plot.

One way to see this is to colour each of the points we draw in Fig. S1a according to the abundance of the species that we drew the point from, with darker points having higher abundance. In METE, we wouldn't expect to see any trends in abundance because any individual draw doesn't depend on the species abundance. However, with the binomial distribution, we would expect that there are very few points with high or low fractions where the species have high abundance. That is, if we have a higher $n_0$, the fraction should always be very close to 0.5 and we should see a trend where darker points cluster to the middle of the curve. The results are shown in Fig. S1b. The individual points are kept in the same order as in Fig. S1a, but offset to better show trends in abundance. We see that the data does not have the same trend as the random placement model, which is why we get a smaller $\alpha$ by maximizing log-likelihood compared to fitting the rank ordering by eye.

In summary, Fig. S1a obscures the abundance data as it only presents the fractions, and the rank ordering hides the trends in abundance that our distributions predict and that we can see in Fig. S1b.

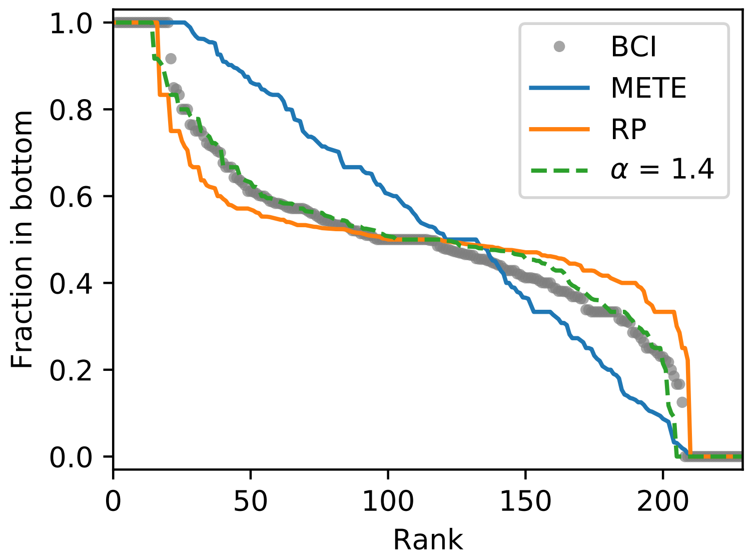


**(a)** Rank ordered

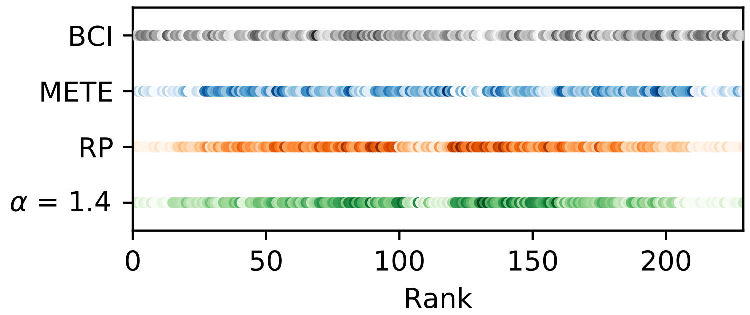


**(b)** Rank ordered, coloured by $n_0$.

**Figure S1.** Rank ordered fraction comparison of METE, random placement, and our density dependent model. Figure S1a seems to indicate that our model is a much better fit. In Fig. S1b the points are coloured according to the log of the abundance with darker points having higher abundance. The points are still in fraction rank order as in Fig. S1a, but offset to better show the trends in abundance. In METE and in the BCI data itself, there isn't much of a trend in abundance, however the binomial distribution and the density dependent case have a clear trend where the most abundant points are clustered in the middle of the distribution. Figure S1a obscures this trend.

| Model | Log-likelihood |
|---|---|
| METE | -729 |
| RP | -963 |
| $\alpha = 1.4$ | -718 |
| $\alpha = 1.12$ | -660 |

**Table S1.** Log-likelihood values for the BCI data set for the three different models, including the by eye fit to the rank ordered plot and the maximum likelihood estimate of $\alpha$. Despite the good apparent fit in the rank ordered plot, here we see that a maximum likelihood approach prefers a smaller $\alpha$.

#### S3 ERROR IN RECOVERING $\alpha$

To test how well we could fit for a known $\alpha$ using our methods, we directly simulate the placement and death processes in Eqs. 4 and 6.

In our simulations, we start with $n_0 = 40$ and then assign a random initial $n$. We simulate 500 placements and deaths with $\alpha = 1.2$ and take the final $n$, repeating this $p$ times to simulate the number of observed points (e.g. species at a single bisection). We then obtain the maximum likelihood $\alpha$.

Figure S2 shows the results of these simulations. Figure S2a shows explicitly how the error scales with the number of points $p$, and Fig. S2b shows the mean $\alpha$ we recover with standard deviations from 5 runs at each number of points.

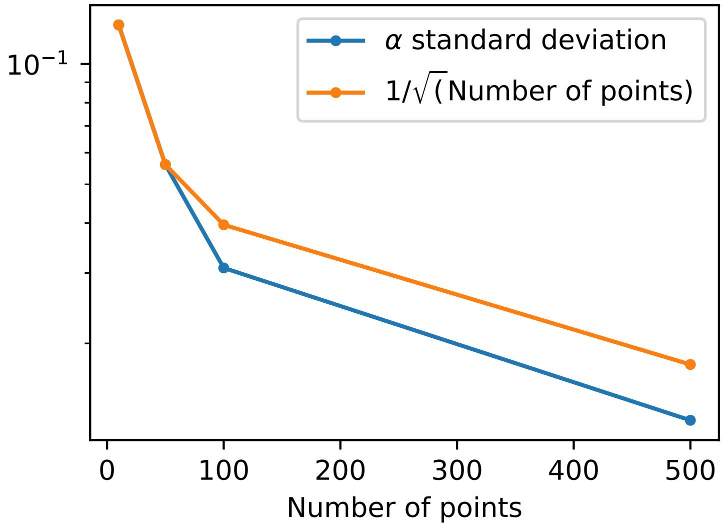


**(a)**

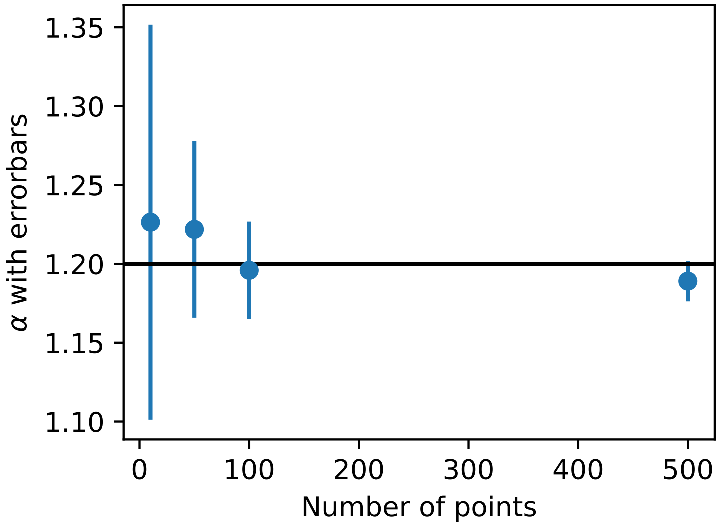


**(b)**

**Figure S2.** The error in recovering $\alpha$ from our simulation scales as 1 over the square root of the number of points $p$. This is shown explicitly in Fig. S2a. Figure S2b shows the value and standard deviation of $\alpha$ that we recover from a simulation with a known $\alpha = 1.2$ (the thick black line). The number of points on the x-axis is analogous to the number of data points we observe in data, and the standard deviation is obtained from doing 5 simulations with that number of runs. We simulated with $p = \{10, 50, 100, 500\}$.

#### S4 RELATING TO THE BINOMIAL DISTRIBUTION

One way to understand why Eq. 9 approaches the binomial distribution for large $n_0$ is to rewrite Eq. 9 as:

$$\Pi_{\alpha=2}(n) = \left( \frac{4(n-n_0/2)^2}{n_0(n_0+1)} + \frac{n_0}{n_0+1} \right) \mathrm{BD}(n, n_0, 1/2). \tag{S6}$$

The first term in the parentheses is 0 at $n = n_0/2$ and approximately 1 ($\frac{n_0}{n_0+1} \approx 1$) at $n = \{0, n_0\}$, and the second term is approximately 1. This means that there is a factor of 2 increase compared to the binomial in the tails of the distribution near $n = \{0, n_0\}$, but around the center the two distributions look identical. Since the tails are already so small at large $n_0$, this factor of 2 isn't really noticeable and so this distribution looks very similar to the binomial distribution.

### S5 RELATING TO THE NEGATIVE BINOMIAL DISTRIBUTION

The negative binomial distribution is very common in ecological modelling (Bliss and Fisher, 1953; He and Gaston, 2000, 2003), and many ecologists are familiar with the $k$ parameter as a measure of aggregation. A standard way to write the negative binomial distribution is

$$\mathrm{NB}(n|p,k) = (1-p)^n p^k \frac{\Gamma(n+k)}{\Gamma(n+1)\Gamma(k)}, \tag{S7}$$

where $p$ is a parameter that relates to the mean $\mu$ of the distribution as $\mu = k(1-p)/p$.

For a bisection, what we want is the conditional negative binomial distribution, where the negative binomial is conditioned on a total of $n_0$ individuals. Given $n$ individuals on one side of the distribution and total number of individuals $n_0$, the corresponding conditional negative binomial is (Conlisk et al., 2007)

$$\mathrm{CNB}(n|n_0,p,k) = \frac{\Gamma(n_0+1)\Gamma(2k)}{\Gamma^2(k)\Gamma(n_0+2k)} \frac{\Gamma(n+k)\Gamma(n_0-n+k)}{\Gamma(n+1)\Gamma(n_0-n+1)}. \tag{S8}$$

We want to compare this to our distribution, Eq. 7. However, we cannot compare directly as the unknown normalization $C$ depends on $\alpha$ and $n_0$. Instead let's look only at the $n$ dependence. If we equate it to the $\Pi$ distribution:

$$\mathrm{CNB} = \frac{\Gamma(n+k)\Gamma(n_0-n+k)}{\Gamma(n+1)\Gamma(n_0-n+1)} \sim \frac{n^\alpha + (n_0-n)^\alpha}{(\Gamma(n+1)\Gamma(n_0-n+1))^{\alpha-1}} = \Pi.$$

Then we want to solve

$$\Gamma(n+k)\Gamma(n_0-n+k) \sim (n^\alpha + (n_0-n)^\alpha)(\Gamma(n+1)\Gamma(n_0-n+1))^{2-\alpha}. \tag{S9}$$

We can directly consider the limiting cases of METE and the binomial distribution from this expression. METE corresponds to $\alpha = 1$, and for the same $n$ dependence we need $k = 1$, which also corresponds to METE. For $\alpha = 2$, which corresponds roughly to the binomial distribution if we ignore the first term on the RHS (good up to a factor of 2, see Eq. S6), the $n$ dependence on the RHS drops out. For this to be true on the LHS, we need $k \to \infty$, which once again is the correct interpretation in terms of aggregation.

It is analytically challenging to equate the $n$ dependence between these two forms more generally, however by plotting the two distributions (Fig. S3) we can see that they are quite similar for specific $k$ and $\alpha$ values.

We can make some comparison by equating the ratios between two different points $n$ and $n'$, eliminating the normalization problem. This is not ideal because we are now only equating at two artificial points, and it is possible the distributions are different outside of that. However, by plotting the distributions (Fig. S3) we can see that matching the peaks of the distributions roughly makes the distributions match at large enough $n_0$, so we will use this approximation.

Taking the ratio gives

$$\frac{\Gamma(n'+k)\Gamma(n_0-n'+k)}{\Gamma(n+k)\Gamma(n_0-n+k)} = \frac{(n'^{\alpha}+(n_0-n')^{\alpha})(\Gamma(n'+1)\Gamma(n_0-n'+1))^{2-\alpha}}{(n^{\alpha}+(n_0-n)^{\alpha})(\Gamma(n+1)\Gamma(n_0-n+1))^{2-\alpha}}.$$

We now equate the ratios of the central points by letting $n' \to n_0/2+1$, and $n \to n_0/2$. This gives

$$\frac{n_0/2+k}{n_0/2+k-1} = \left(1+\frac{2}{n_0}\right)^{2-\alpha}\left(\frac{1}{2}\left(1+\frac{2}{n_0}\right)^{\alpha}+\frac{1}{2}\left(1-\frac{2}{n_0}\right)^{\alpha}\right).$$

Expanding around large $n_0$ and keeping the first order term gives

$$k \approx \frac{n_0}{2}\left(\frac{\alpha-1}{2-\alpha}\right).$$

For $\alpha = 1$, we know $k = 1$, so we use that as the constant offset to get our approximate relationship

$$k \approx \frac{n_0}{2}\left(\frac{\alpha-1}{2-\alpha}\right)+1. \quad \text{(S10)}$$

Figure S3 plots the conditional negative binomial distribution with $k$ calculated from Eq. S10 compared to our distribution for a range of $\alpha$s. Our derived approximate relationship results in good agreement between these distributions.

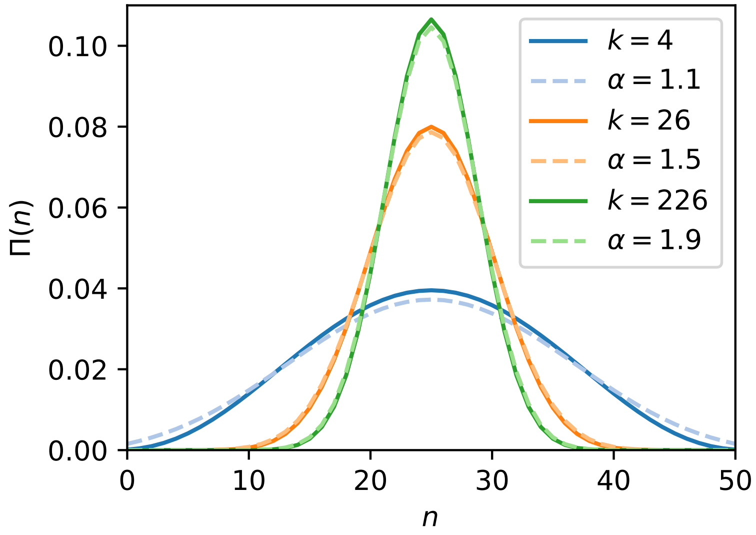


**Figure S3.** Conditional negative binomial distributions (solid lines) with $k$ calculated from Eq. S10 compared to our predicted distributions with their corresponding $\alpha$ (dashed lines). $n_0 = 50$.

#### S6 TRENDS IN DIAMETER AND ABUNDANCE FOR BCI DATA

Figure S4 shows the relationship between $\alpha$ and species abundance at different scales, using the BCI data. At smaller scales, the distribution of $\alpha$ is broader and generally higher, as we found in Fig. 2. There aren't any high abundance high $\alpha$ values.

Figure S5 shows the relationship between $\alpha$ and the mean dbh for each species at different scales, again using the BCI data. We still see the trend that at smaller scales the $\alpha$ distribution is broader, but at no scale is there an apparent relationship between $\alpha$ and the mean dbh. We might expect that larger species have higher $\alpha$ values at larger scales than smaller species, as they are competing at a different scale, however we don't see that effect. It is possible that bisections are not sufficient to pick up on this difference, or that intraspecies size variation obscures this trend. Additionally, the range of species' mean dbh here is only from about 100 mm to about 500 mm, so it is also possible that we need to consider a much larger range of dbh before we see this effect. Finally, as mentioned in the main text, it is possible we did not go to scales small enough to see the difference in aggregation from size variation.

Finally, we can test if the lack of high $\alpha$ high $n_0$ species is a consequence of energy equivalence. That is, the most abundant species are also the smallest. We didn't see a trend in dbh alone, but we may see a trend in species total metabolic rate. For trees, metabolic rate scales approximately as dbh$^2$. Figure S6 shows how $\alpha$ scales with the total metabolic rate of all individuals in a species (which scales as $n_0 \times \text{dbh}^2$). Again, we see no significant trend.

That we see no trends in any of these plots might indicate that the operating density dependent mechanism is not resource limitation. One plausible mechanism is the Janzen-Connell effect.

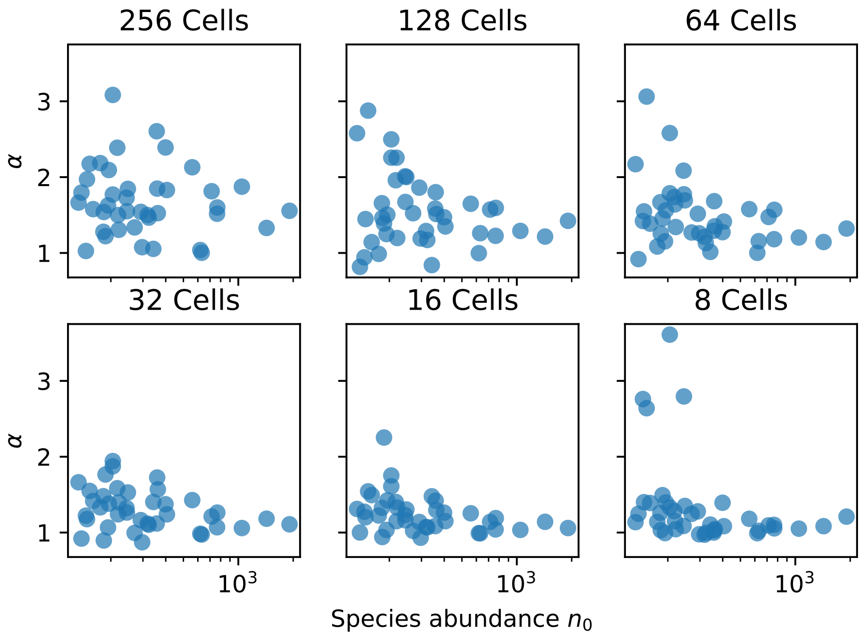


**Figure S4.** Species abundance and their corresponding maximum likelihood $\alpha$ value using the BCI data at a range of spatial scales from 3 bisections to 8 bisections. The absolute scale is 50 ha divided by the number of cells.

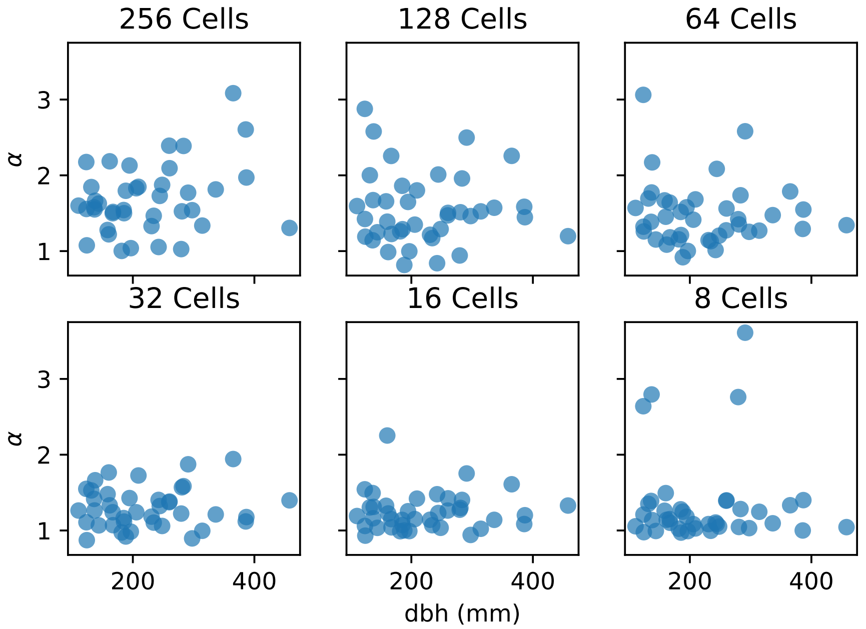


**Figure S5.** Mean dbh in mm for each species and their corresponding maximum likelihood $\alpha$ value using the BCI data at a range of spatial scales from 3 bisections to 8 bisections. The absolute scale is 50 ha divided by the number of cells.

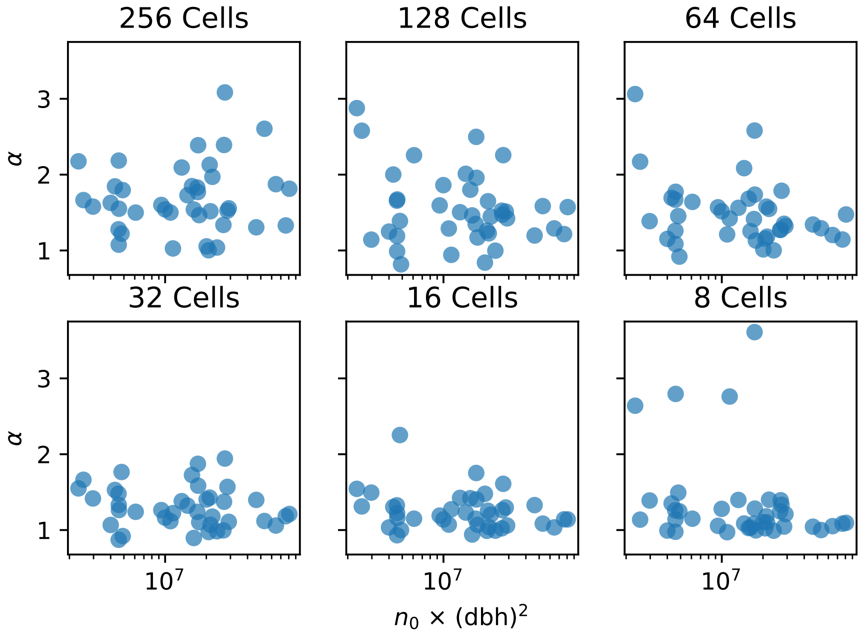


**Figure S6.** Each species' total metabolic rate scales as $n_0 \times \mathrm{dbh}^2$, which is plotted against their corresponding maximum likelihood $\alpha$ value using the BCI data at a range of spatial scales from 3 bisections to 8 bisections. The absolute scale is 50 ha divided by the number of cells.
